# In-Silico Evaluation of Silk Fibroin-Conjugated Doxycycline for Sustained Drug Delivery

**DOI:** 10.1101/2025.07.31.668034

**Authors:** Kranti Kiran Reddy Ealla, Sreenivasan Kuravi, Vikas Sahu, Chandra Sri Durga, Neema Kumari, Kiran Kumar Bokara

## Abstract

Tetracyclines such as Doxycycline and Minocycline are broad-spectrum antibiotics. Their high oral bioavailability aids systemic absorption but limits achieving sustained local drug levels at the infection site. Sustained drug release to enhance the therapeutic efficacy and minimise side effects remains a significant challenge. This study investigated the conjugation of Doxycycline with silk fibroin (SF), which would provide sustained drug release at the target site. We employed an in-silico approach using molecular docking and molecular dynamics (MD) simulations to investigate the interaction and stability of antibiotics with SF. We also assessed the structural stability of SF when conjugated with these antibiotics. Molecular level characterization using a blind docking approach and MD simulation revealed that Doxycycline exhibited the strongest binding affinity of -6.0 kcal/mol, compared with both Tetracycline (−5.9 kcal/mol) and Minocycline (−5.7 kcal/mol). The MMPBSA analysis also showed that Doxycycline exhibited the highest binding energy of -25.71 kcal/mol compared to Tetracycline (−18.27 kcal/mol) and Minocycline (−10.8 kcal/mol). The strong binding energy and stability of Doxycycline with SF was attributed mainly by van der Waals interactions and hydrogen bonds. To evaluate the stability of protein-antibiotic complexes, 100 ns MD simulations were performed. The RMSD, RMSF, radius of gyration, SASA, and DSSP analyses revealed that SF was stable without much deformation upon docking of all three antibiotics. In summary, our computational results suggest that Doxycycline and SF conjugates offer a promising strategy for sustained drug delivery, potentially improving therapeutic efficacy in the treatment of chronic inflammatory diseases.

**Highlights:**

- The study finds silk fibroin as a potential carrier for sustained drug delivery of tetracycline family antibiotics.
- Silk fibroin remained stable with minimal structural deviations in complex with Doxycycline, Minocycline, and Tetracycline over a 100 ns MD simulation.
- Doxycycline showed the highest binding affinity and most favourable binding energy and remained stable throughout the simulation.
- SF-Doxycycline conjugate shows promise for sustained, localized drug delivery.

## 1.0 Introduction

Tetracyclines are broad-spectrum antibiotics with antimicrobial activity(Klein & Cunha, 1995). The clinical efficacy of these antibiotics has already been reported in various dermatological diseases, autoimmune disorders, and corneal inflammation, and is widely used in dentistry (Sapadin & Fleischmajer, 2006). Doxycycline and Minocycline are second-generation tetracyclines having a broad spectrum of antimicrobial effects and pharmacokinetic properties, mostly employed for treating various inflammatory diseases, oral infections such as gingivitis and periodontitis (Khatri & Kumar, 2012; Klein & Cunha, 1995; Spasovski et al., 2016). These antibiotics exhibit high oral bioavailability (90–100%) and are lipophilic, allowing them to cross cell membranes and dissolve rapidly in body fluids(Agwuh & MacGowan, 2006). However, this systemic diffusion may reduce their therapeutic effect at the site of infection, making the treatment less effective. Therefore, conventional dosages are employed for a prolonged period for effective results at the target site in periodontal treatment. High dosage administration for a prolonged period leads to adverse side effects(Majewski, 2014), and a sustained drug release system needs to be employed for effective treatment and to increase therapeutic efficacy and decrease treatment time.

Developing biocompatible and efficient drug delivery systems is a critical area of research in modern medicine. Systemic toxicity, poor bioavailability, and a lack of targeted delivery are general problems with traditional drug administration. Controlled release systems improve therapeutic efficacy and minimize adverse side effects due to conventional dosage, and maintain optimal drug concentrations(Adepu & Ramakrishna, 2021). A drug delivery carrier should possess biocompatibility, biodegradability, thermal stability, a high surface to charge ratio to increase the pharmacokinetics of the drug for efficient loading and a controlled release mechanism(Bandopadhyay et al., 2019; Finbloom et al., 2020). Natural biopolymers gained great attention in drug delivery, among them, Silk fibroin emerged as a promising biomaterial due to its biocompatibility, non-immunogenic and non-antigenic nature, biodegradation, strength, and stability due to its β-sheet backbone conformation. These properties were validated in various in vivo models, establishing SF as a good drug delivery carrier(Wenk et al., 2011).

Silk fibroin is a natural biomaterial extracted from the silk worm cocoons and has been investigated as a drug delivery system for various small molecules and drugs(Pritchard & Kaplan, 2011). Silk fibroin has been used as a carrier for sustained drug delivery of amphiphilic drugs such as curcumin and emodin(Montalbán et al., 2020a). Silk-based drug delivery systems were reported for sustained drug release of various anticancer drugs such as Paclitaxel, Doxorubicin, Curcumin, Methotrexate, and Emodin(Jastrzebska et al., 2015). Silk fibroin was used as a carrier of antibiotics to mitigate the burst release and increase the bioavailability at the target site. Antibiotics such as Vancomycin, Gentamycin, Tetracycline, Colistin, Amoxicillin trihydrate, and Ciprofloxacin hydrochloride conjugated with silk fibroin for various clinical applications such as wound healing, bone tissue engineering, treatment of osteomyelitis, and development of sutures, ear tubes, and orthopedic implants(Han et al., 2017)(Ghalei & Handa, 2022). Antibiotic loaded silk fibroin sutures were employed in wound suturing to enable sustained antibiotic release and enhance antimicrobial efficacy(Choudhury et al., 2016). Silk fibroin-coated Azithromycin microspheres showed sustained release of antibiotic at the target site and ameliorated inflammation, and induced periodontal tissue regeneration(Ouyang et al., 2025).

Doxycycline has been incorporated into various biomaterial matrices such as chitosan, collagen, gelatin, hyaluronic acid in the form of microspheres, nanoparticles, nano-fibrous membranes, and hydrogels, for various biomedical applications including periodontosis(Gjoseva et al., 2018; Stan et al., 2023; Toledano-Osorio et al., 2023). However, silk fibroin and Doxycycline conjugates remain largely unexplored. In our previous studies, Doxycycline incorporated SF hydrogels exhibited anti-bacterial activity showed sustained release, and showed a lesser zone of inhibition than Doxycycline alone(Ealla et al., 2025). Though tetracycline family drugs were conjugated with silk-based formulations in various clinical applications, their interaction and stability, and the stability of the carrier (silk fibroin) through computational studies have not yet been reported. Our present study investigated the interaction profile of tetracycline family drugs Doxycycline, Tetracycline, and Minocycline with Silk fibroin matrix by molecular docking, and to evaluate the stability of both protein and protein-drug complex by employing a computational approach of molecular docking analysis and molecular dynamic simulations.

## 2.0 Materials and Methods

### 2.1 Data sets

The crystal structure of silk fibroin (PDB ID: 3UA0) was retrieved from the Protein Data Bank (PDB). The 3D structures of the ligands Doxycycline hyclate (PubChem CID 54686183), Minocycline (PubChem CID 54675783), and Tetracycline (PubChem CID 54675776) were obtained from the PubChem database.

### 2.2 Visualisation and plotting tools

Molecular visualising tools such as PyMOL were used to create 3D interaction profiles of protein-ligand complexes, and UCSF Chimera(Pettersen et al., 2004) is used to create superimposed images. LigPlot + v.2.2(Laskowski & Swindells, 2011; Wallace et al., 1995) is used to create 2D interaction plots of Protein-ligand complexes, to demonstrate the hydrogen and hydrophobic interactions between the ligand and residues of the protein.

### 2.3 Molecular Docking

Molecular docking was performed to determine the interaction and binding affinity of Doxycycline, Tetracycline, and Minocycline with Silk fibroin protein. Molecular docking and binding energy calculations were performed using AutoDock Vina(Eberhardt et al., 2021). The protein and ligands were prepared for docking using AutoDock Tools 1.5.7. Water molecules were removed, Kollman charges and polar hydrogen were added to the protein, and Gasteiger charges were added to the ligands. A blind docking protocol was followed, and a grid box size with dimensions x = 126 Å, y =126 Å, z = 126 Å in each direction was prepared to encompass the entire protein, ensuring the docking search space covers all possible binding sites. The molecular docking assessment was carried out using command-line (CMD) instructions for AutoDock vina(Montalbán et al., 2020b)(Behera et al., 2023). The results were visualized using PyMOL, Discovery Studio, ChimeraX, and LigPlot+.

### 2.4 MD Simulations and Analysis

Molecular dynamics (MD) simulations were performed using GROMACS 2021 to analyze ligand-induced conformational changes and dynamic behavior at the atomic level(Berendsen et al., 1995). The top-scoring pose from blind docking served as the initial protein-ligand conformation. The protein topology file was generated using the pdb2gmx tool, and ligand topology files were generated using CHARMM-GUI(Brooks et al., 2009), and the complexes were solvated in a triclinic box with the simple point charge (SPC) water model. System neutrality was maintained using 0.1 M NaCl. The CHARMM27 force field was applied, and energy minimization was conducted using the steepest descent and conjugate gradient algorithms for a maximum of 50,000 steps. Equilibration was carried out under NVT and NPT conditions at a temperature of 300 K and a pressure of 1 bar. Finally, the production phase of simulations was carried out for 100 ns (10000 ps) for each system. The resultant trajectory files after the simulation were analyzed using GROMACS tools to calculate Root Mean Square deviation, Root Mean Square fluctuations, Radius of gyration, Solvent Accessible Surface Area, and hydrogen bonds (Berendsen et al., 1995). The data analysis and plots were generated using Grace software (https://plasma-gate.weizmann.ac.il/Grace/).

### 2.5 MM-PBSA binding free energy calculations

Binding free energy was calculated using the Molecular Mechanics Poisson-Boltzmann Surface Area (MMPBSA) method implemented using the gmmpbsa tool available in the GROMACS package. The Python script, MmPbSaStat.py, available in the g_mmpbsa package, was used to calculate the average binding free energies of all the protein-ligand complexes. The binding free energy of the complexes is computed based on the prediction of solvation and molecular mechanics potential energies(Kumari et al., 2014).

### 2.6 Superimposition analysis

The superimposition analysis was performed to identify the structural changes in the Silk fibroin protein structure after a 100 ns simulation. The Matchmaker program of UCSF Chimera X was used for superimposition by employing the Needleman-Wunsch alignment algorithm and BLOSUM-62 similarity matrix(Meng et al., 2006).

## 3. Results and discussion

### 3.1 Molecular Docking

Molecular docking serves as a pivotal computational technique in structural biology and drug discovery, aiming to anticipate the optimal orientation of a ligand as it binds to a larger biomolecule (receptor), forming a stable complex. In SF, specific binding sites have not been experimentally characterised, so blind docking was employed for unbiased exploration of ligand binding sites on the entire molecular surface of the protein(Hetényi & Van Der Spoel, 2006; Montalbán et al., 2020b). The three ligands, Doxycycline, Tetracycline, and Minocycline, displayed better docking scores with SF, and all three ligands interacted with the residues of both A and B chains of Silk Fibroin. The top three affinity scores were selected from the Vina output log files, and the average of those top three scores was considered as the predicted binding affinity scores and plotted in a scatter plot, respectively, as shown in Figure 1.

**Figure 1:**
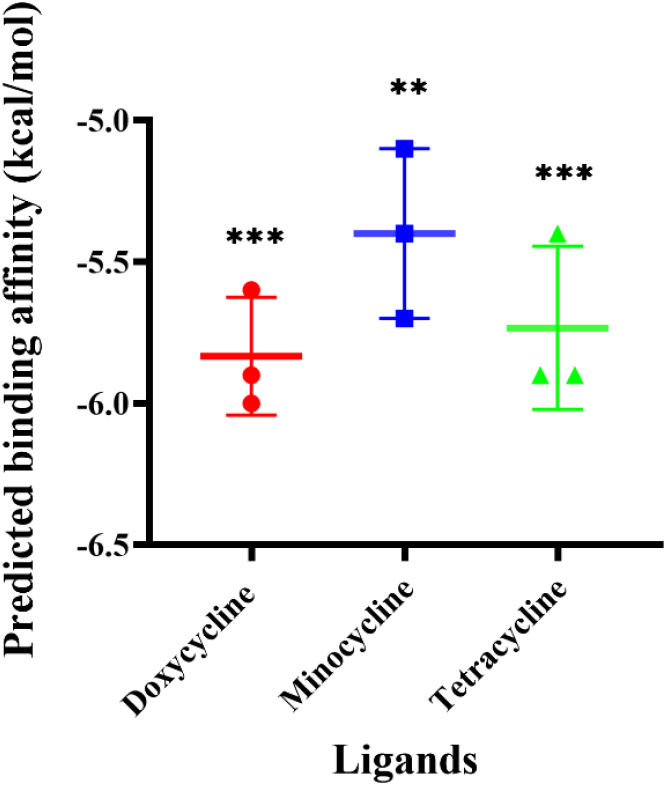
Scatter plot showing predicted binding affinity (kcal/mol) of ligands Doxycycline, Minocycline, and Tetracycline against silk fibroin protein.

Doxycycline, Tetracycline, and Minocycline displayed a significant docking score of -6.0 Kcal/mol, 5.9 kcal/mol, and 5.7 Kcal/mol, respectively. Among the three ligands, Doxycycline showed the highest binding affinity, and the lowest binding affinity was displaced by Minocycline. All three ligands formed hydrogen and hydrophobic interactions. The interaction profiles of the ligands with SF are depicted in Table 1. The Doxycycline formed three hydrogen bonds, one with Asn(B)68 and two with Glu(A)94, respectively. Tetracycline formed four hydrogen bonds with residues, one each with Glu(A)28, Tyr(A)30, Phe(A)31, and Val(A)37. Minocycline formed one hydrogen bond with Arg(A)48. The molecular interaction profiles, such as the 2D (LigPlot interaction), and the complete surface illustration of the protein-ligand complexes were depicted as in Figure 2.

**Table 1:** Molecular docking results showing docking score, number of hydrogen bonds formed, amino acids of silk fibroin involved in hydrogen and hydrophobic interactions with ligands Doxycycline, Minocycline, and Tetracycline.

| Sl.No | Drug/ ligand | Docking score (Kcal/mol) | Number of H-bonds | Residues of SF involved in H-bonding | Residues of SF involved in hydrophobic interactions |
| --- | --- | --- | --- | --- | --- |
| 1 | Doxycycline | 6.0 | 3 | Glu(A)94, Asn(B)68 | Ile(A)86, Ile(B)23, Asp(B)25, Phe(B)26, Gln(B)65, Lys82(B) |
| 2 | Minocycline | 5.7 | 2 | Arg(A)48 | Ile(A)46, Val(A)54, Ser(B)107, Asp(B)108 |
| 3 | Tetracycline | 5.9 | 4 | Glu(A)28, Tyr(A)30, Phe(A)31, Val(A)37 | Asp(A)29, Thr(A)36, Gln(A)38 |

**Figure 2:**
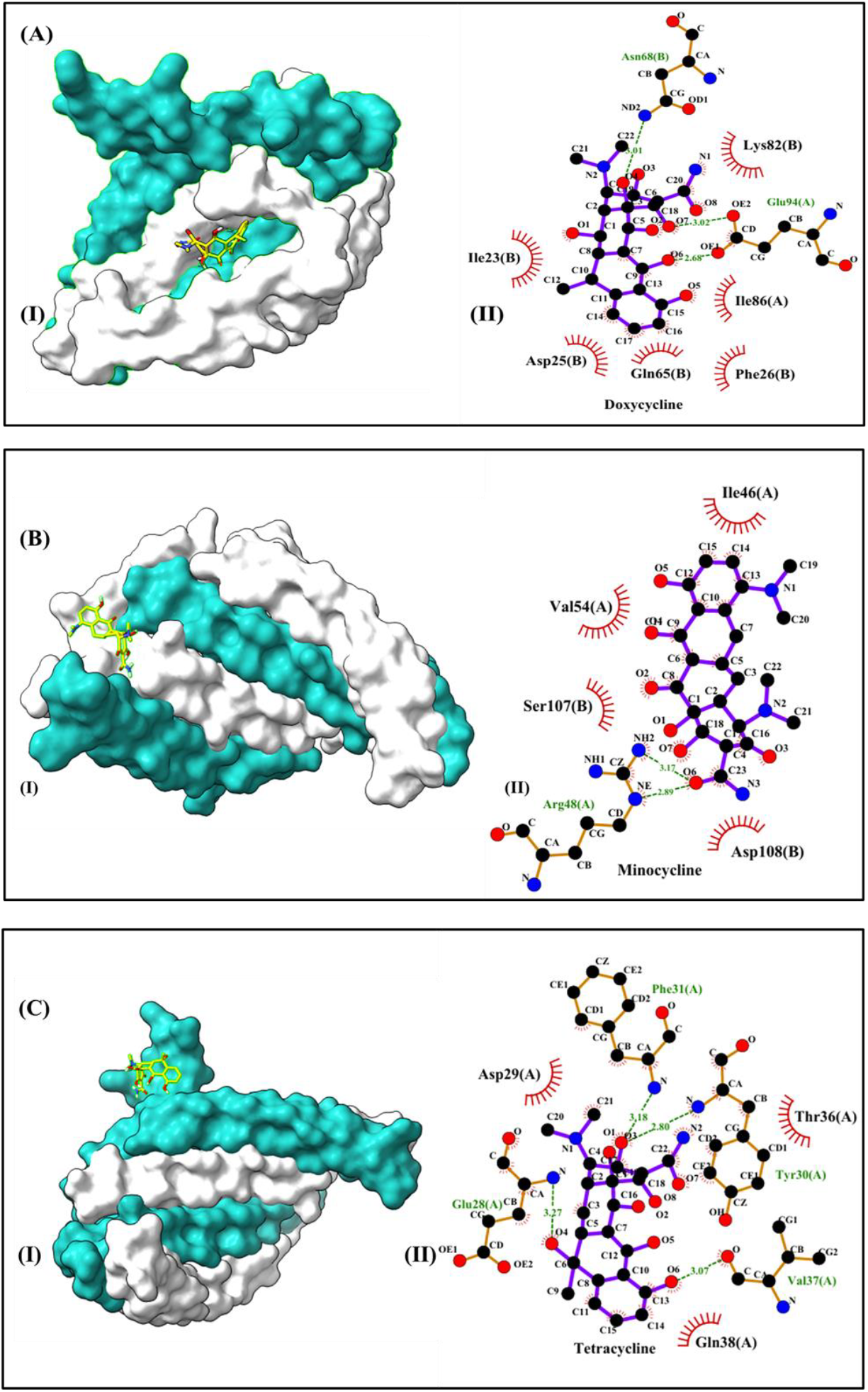
Molecular binding and interaction profiles of SF-ligand complexes (A) Doxycycline, (B) Minocycline, (C) Tetracycline, showing (I) Complete surface illustration of the SF structure showing the ligand-binding interface, with the A-chain in cyan, B-chain in white, and ligands represented in yellow. (b) 2D LigPlot interaction diagrams illustrating hydrogen bonds (shown in green) and hydrophobic interactions between the ligand and silk fibroin (SF).

### 3.2 Molecular dynamics simulations

Molecular dynamics simulation is a widely accepted technique to comprehensively investigate the stability of protein-ligand interactions(De Vivo et al., 2016). Over the past decade, MD simulations have emerged as an important tool for studying the drug delivery applications, such as carrier-drug interactions, carrier drug stability, and drug solubility(Katiyar & Jha, 2018). To analyze the structural and conformational changes, such as the stability and dynamic characteristics of protein-ligand complexes, a molecular dynamics simulation (MDS) was carried out for 100 ns. The structural alterations and flexibility of all ligand-bound protein complexes were determined using various stability parameters that include backbone and ligand Root-Mean-Square Deviations (RMSD), Root-Mean-Square Fluctuations (RMSFs), the Radius of gyration (Rg), and Solvent Accessible Surface Area (SASA) during the 100 ns simulation.

#### 3.2.1 Root Mean Square Deviation

Root Mean Square Deviation (RMSD) is a widely used metric in molecular dynamics (MD) simulations to measure the structural deviation of a protein bound to a ligand over time relative to a reference structure(Sargsyan et al., 2017). The conformational and structural variations in the backbone atoms of SF and SF-ligand complexes were studied using RMSD analysis depicted in Figure 3A. The unbound SF exhibited an initial RMSD of approximately 0.55 nm – 0.6 nm from 10 ns to 42 ns, gradually increasing to 0.63 nm at 42 ns and fluctuating between 0.60–0.70 nm until 52 ns. Further, the protein fluctuated between 0.5-0.65 nm till the end of the simulation. The protein in the presence of all three drugs showed lower RMSD compared to the protein alone. The RMSD of the SF-Doxycycline complex remained within 0.35–0.45 nm from 10 ns to 40 ns, followed by a gradual increase to 0.45–0.60 nm up to 63 ns. Notably, from 63 ns to 100 ns, the complex stabilized further, with RMSD values consistently ranging between 0.40 and 0.55 nm. SF-minocycline complex stabilised at an RMSD range of 0.37nm till 20ns of simulation further and fluctuated within in range of 0.3 – 0.5nm till 40ns and stabilised at 0.37 nm till the end of simulation. The RMSD of the SF-Tetracycline complex gradually increased from 0.25 nm to 0.5 nm till 40 ns of simulation and fluctuated around 0.4 nm till 90 ns, and higher fluctuations were observed than SF alone at the end of the simulation. Among the three ligands, Doxycycline showed a stable and stronger interaction than Minocycline and Tetracycline with the least RMSD of 0.05 nm as shown in Figure 3B.

**Figure 3:**
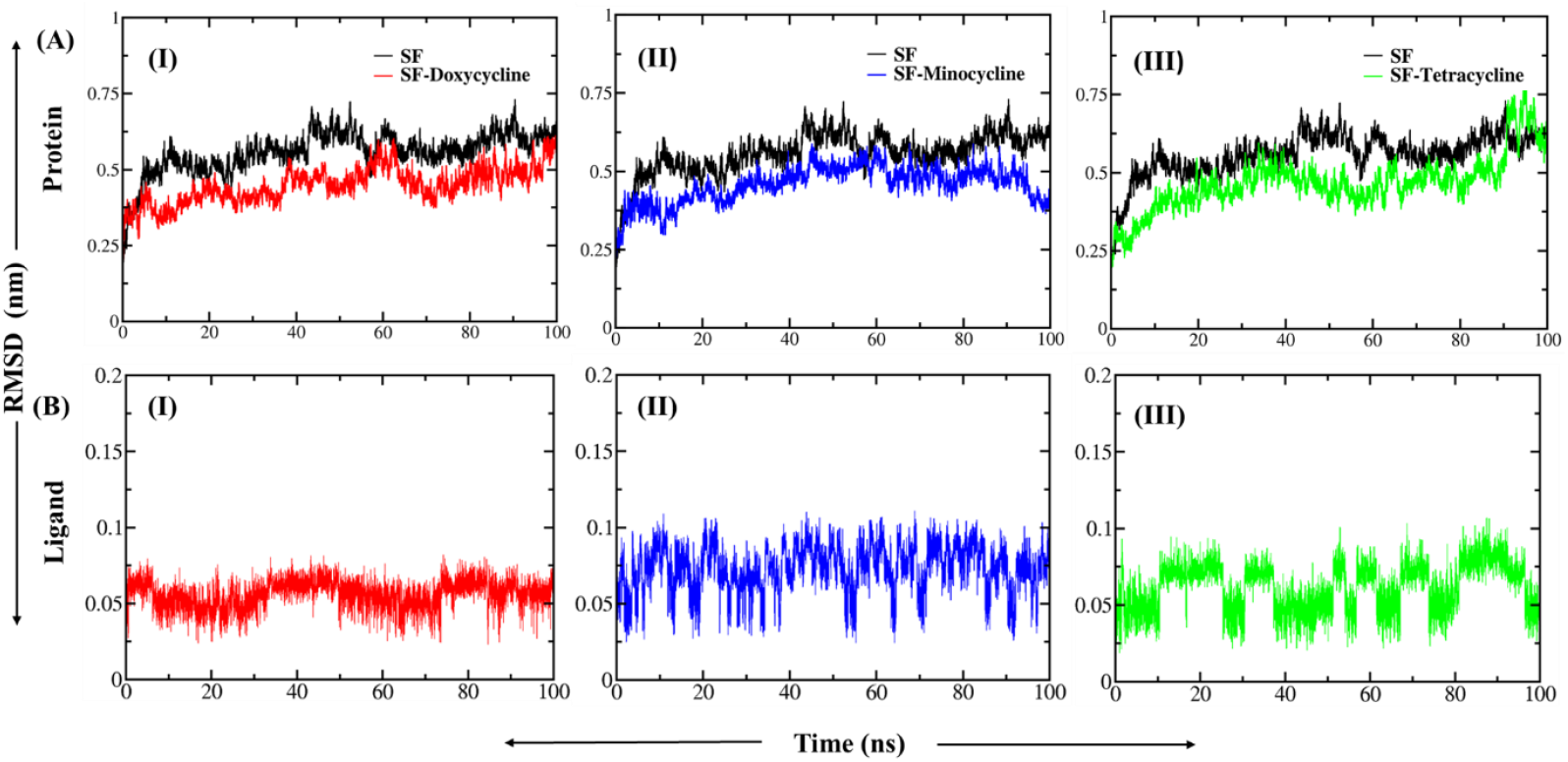
**A)** The RMSD pattern of SF complexed with ligands. **(I)** RMSD of SF-Doxycycline complex (coloured red) compared with SF alone (coloured black), **(II)** RMSD of SF-Tetracycline complex (coloured green) compared with SF, **(III)** RMSD of SF-Minocycline complex (coloured blue) compared with SF. **(B)** RMSD of ligands complexed with SF.**(I)** Doxycycline **(II)** Minocycline, **(III)** Tetracycline.

#### 3.2.2 Root Mean Square Fluctuations (RMSF)

Root mean square fluctuation analysis was performed to measure residue-wise fluctuations in SF and SF-ligand complexes. The RMSF method effectively quantifies the flexibility of the protein by measuring the displacement of individual amino acids from their average position during simulation. The high RMSF values in the protein indicate flexibility regions with coils, turns, and loops, whereas low RMSF values indicate stable and rigid regions of the protein with alpha helices and beta sheets as secondary structure(Martínez, 2015). Silk fibroin consists majority of beta sheet and coils as secondary structures in its backbone. The RMSF plots for SF and SF-ligand complexes were depicted in Figure 4. The PDB structure of SF contains identical A-chain and B-chain. The A-chain contains residues 26-108, and the B-chain contains 23-108 residues. The residue numbers in Figure 4 are interpreted according to the PDB (3UA0).

**Figure 4.**
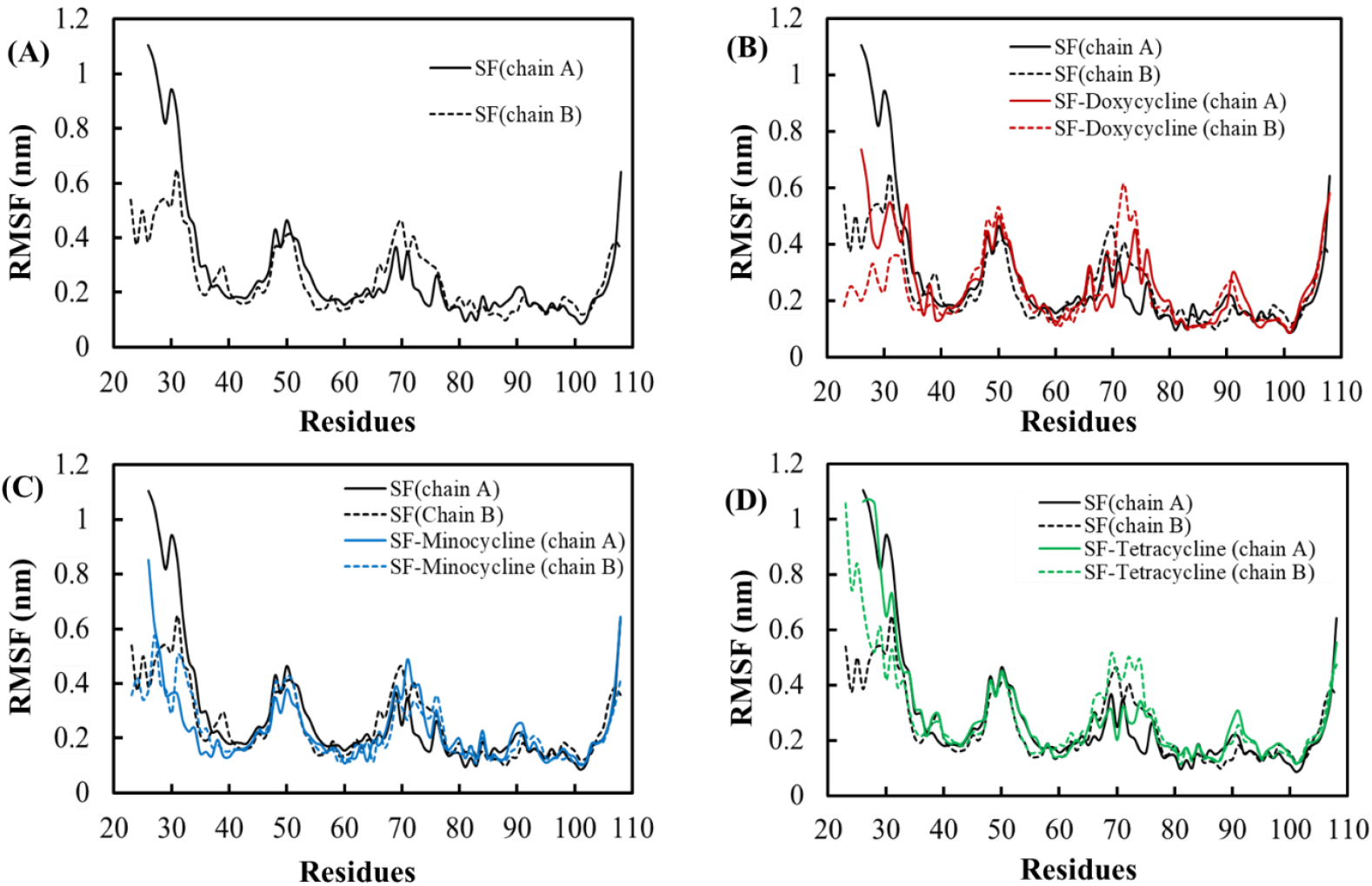
RMSF graphs of SF and SF-ligand complexes showing **(A)** SF alone, **(B)** SF-Doxycycline complex, **(C)** SF-Minocycline complex, **(D)** SF-Tetracycline complex. In each panel, the A-chain is represented by a solid line and the B-chain by a dotted line.

The RMSF plot of SF alone shows an increase in RMSF values of residues at the coil regions, indicating flexibility, whereas residues at the beta sheet region exhibited lower RMSF values. The N-terminal and C-terminal residues of the protein exhibited higher fluctuations, and the core region of the protein remained stable, with notable fluctuations in the coil structures. The SF-Doxycycline complex showed a decrease in the fluctuation in the N-terminal residues of both the A and B chains, and a slight increase in fluctuation was noted around the residues interacting with Doxycycline, as shown in Figure 4. The average RMSF of the SF-Doxycycline complex compared to SF alone was decreased from 0.27 nm to 0.25 nm for the A-chain and 0.25 nm to 0.23 nm for the B-chain, respectively. The RMSF of the SF-Minocycline complex was similar to SF alone, and no major changes were observed except for the decrease in RMSF of N-terminal residues. Overall, the average RMSF of the SF-Minocycline complex compared to SF alone was decreased from 0.27 nm to 0.23 nm for the A-chain and from 0.25 nm to 0.22 nm for the B-chain. The SF-Tetracycline complex showed the RMSF similar to the SF-Doxycycline complex. The N-terminal residues of the B-chain interacted with tetracycline and showed more fluctuations than SF alone. The average RMSF of the SF-Tetracycline complex compared to SF alone was slightly increased from 0.27 nm to 0.28 nm for the A-chain and from 0.25 nm to 0.29 nm for the B-chain. Over all RMSF analysis suggests that all the SF remained stable upon binding of all three ligands throughout the simulation, indicating the stability of the protein, supporting the suitability of SF as a stable drug carrier for Tetracycline-class antibiotics.

#### 3.2.3 Radius of gyration (Rg)

The radius of gyration (Rg) describes the overall compactness of a molecule, such as a protein, nucleic acid, polymer, or protein-ligand complex. Rg serves as a marker of how compact the arrangement of protein is. The higher Rg range implies a decrease in protein structural compactness, an increase in elasticity, and less stability(Lobanov et al., 2008). The plot of Rg as a function of time for Silk Fibroin and all Silk Fibroin-ligand complexes for the 100 ns trajectory was depicted in Figure 5A.

**Figure 5.**
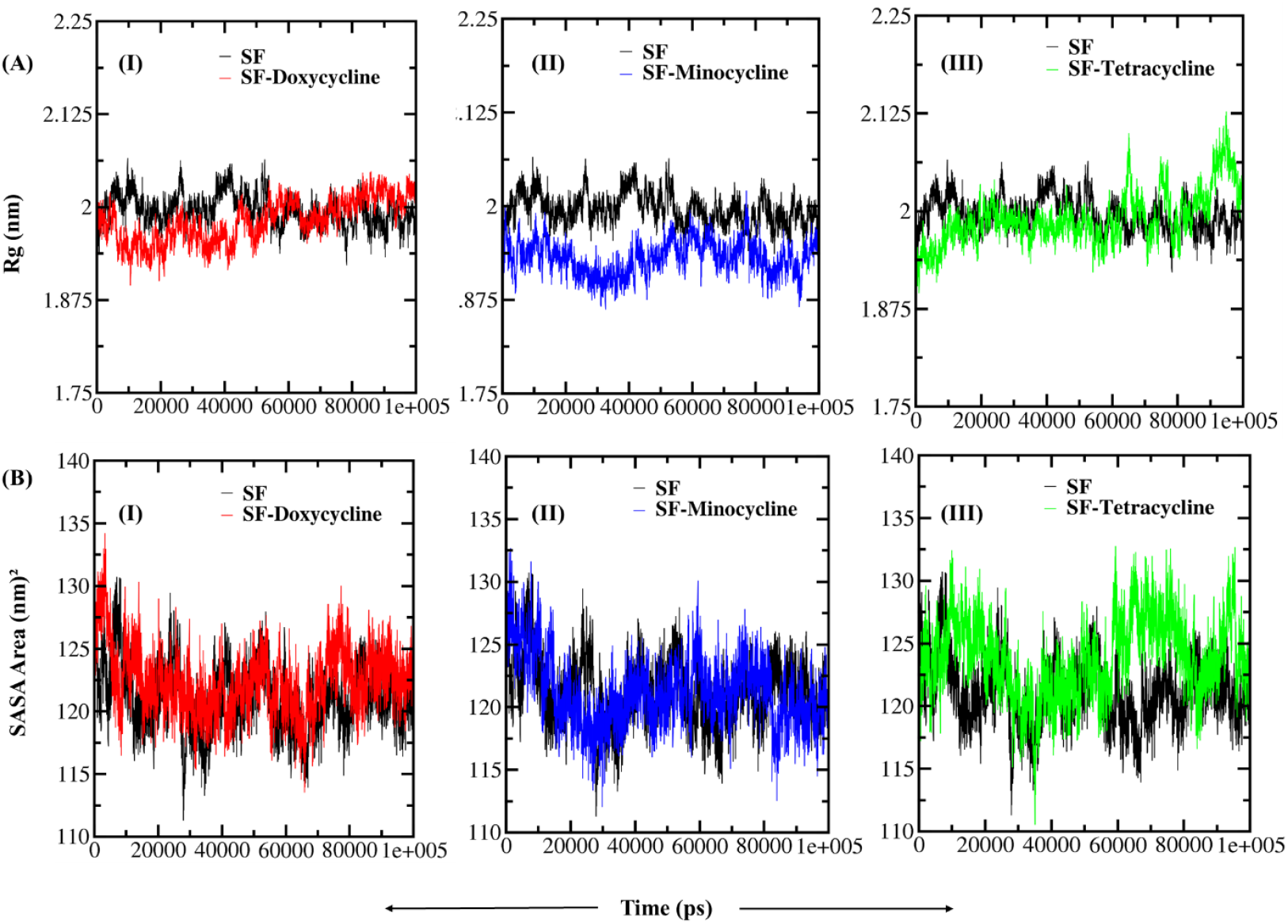
**(A)** Rg as a function of time for SF and SF-ligand complexes over a 100 ns MD simulation. **(I**) Rg of SF-Doxycycline complex (coloured red) **(II)**, Rg of SF-Minocycline complex (coloured blue) **(III)**, Rg of SF-Tetracycline complex (coloured green) compared with the Rg of unbound SF (coloured black). **(B)** SASA as a function of time for SF and SF-ligand complexes over a 100 ns MD simulation. SASA profile of **(I)** SF-Doxycycline complex (coloured red), **(II)** SF-Minocycline complex (coloured blue), **(III**) SF-Tetracycline complex (coloured green) compared with the SASA profile of unbound SF (coloured black).

The SF alone showed Rg around 2 nm and was consistent till the end of the simulation. The Rg value of all protein ligand complexes was computed to be in the range from 1.85 nm to 2.0 nm with varying functions of time. The Rg of the SF-Doxycycline complex was from 1.93 nm to 2 nm and closely aligned with the Rg of SF alone. The SF-Minocycline complex showed Rg lower than SF, ranging from 1.87 nm to 1.93 nm. The SF-Tetracycline complex showed Rg range along the SF till 60-ns and showed an increase in Rg and fluctuations from 60 ns to the end of the simulation. Among the three ligands, Minocycline exerted the most stabilizing effect on SF by maintaining lower and consistent Rg values, indicating a compact and stable complex. Doxycycline provided moderate stabilization, while Tetracycline appeared to reduce the structural integrity of the SF complex over time.

#### 3.2.4 Solvent Accessible Surface Area (SASA)

Solvent accessible surface area (SASA) is described as the area of the protein that is exposed enough to make interactions with the neighbouring solvent molecules(Durham et al., 2009). SASA was performed to determine the extent to which the amino acids of the protein are exposed to solvent. SASA analysis of SF and its complexes with Doxycycline, Minocycline, and Tetracycline was depicted in Figure 5B.

The SF alone showed a SASA range between 120 and 130 nm^²^ till 20 ns of simulation, and further fluctuated between 115 and 127 nm^²^ till 40 ns, and showed a SASA range around 120 nm^²^ from 40 ns till the end of the simulation. The SASA of the SF-Doxycycline complex showed 130 nm^²^ during the start of the simulation and gradually decreased to 117 nm^²^ and was stable till 50 ns, and further fluctuated around 122 nm^²^ till the end of the simulation. The SF-Minocycline complex showed the SASA range of 127 nm^²^ at the start of the simulation and continuously decreased to 117 nm^²^ till 30 ns, and from then on stabilised around 120 nm^²^ till the end of the simulation. The SF-Tetracycline complex showed broader SASA variations throughout 100 ns. The fluctuations observed in the SF-Tetracycline complex suggest a dynamic interaction environment, potentially indicating more significant conformational changes compared to the other complexes. Overall, these variations in SASA highlight the differing stability and behaviour of the complexes under the simulated conditions. The complex showed wider fluctuations between 115 and 132 nm^²^, with no significant stabilisation till the later stages of simulations.

Overall, both ligands, Doxycycline and Minocycline, confer stability and compactness upon binding to the SF protein, whereas the Tetracycline complex demonstrated persistent fluctuations throughout the simulation. Such behaviour may imply less effective encapsulation or stabilization potential when compared to Doxycycline and Minocycline.

#### 3.2.5 Secondary structure profile analysis

The stability of protein varies according to the variations in protein secondary structures. Differences in the secondary structure elements, such as α-helices, β-sheets, coils, and turns, were investigated using the DSSP module of GROMACS to check the conformational dynamics and folding mechanism of SF upon complexing with ligands(Tanwar & George Priya Doss, 2018). The Secondary structure profiles of SF and SF-ligand complexes were depicted in Figure 6. The SF majorly contains β-sheet, the major backbone of its protein structure, inducing stability and strength. It also has coils and turns, indicating minor flexibility in the protein structure. Throughout the simulation, the secondary structure of the SF protein remained stable, suggesting no significant unfolding or disruption in the absence of ligand. Upon binding of Doxycycline and Tetracycline, a slight increase in the coil content and a decrease in β-sheet content were observed, suggesting flexibility and partial unfolding of protein structure upon binding of these ligands. The β-sheet content of Minocycline-bound SF was similar to SF, but a slight increase in coil and turn content was observed, suggesting Minocycline may exert a minimal destabilisation effect on the secondary structure of SF.

**Figure 6.**
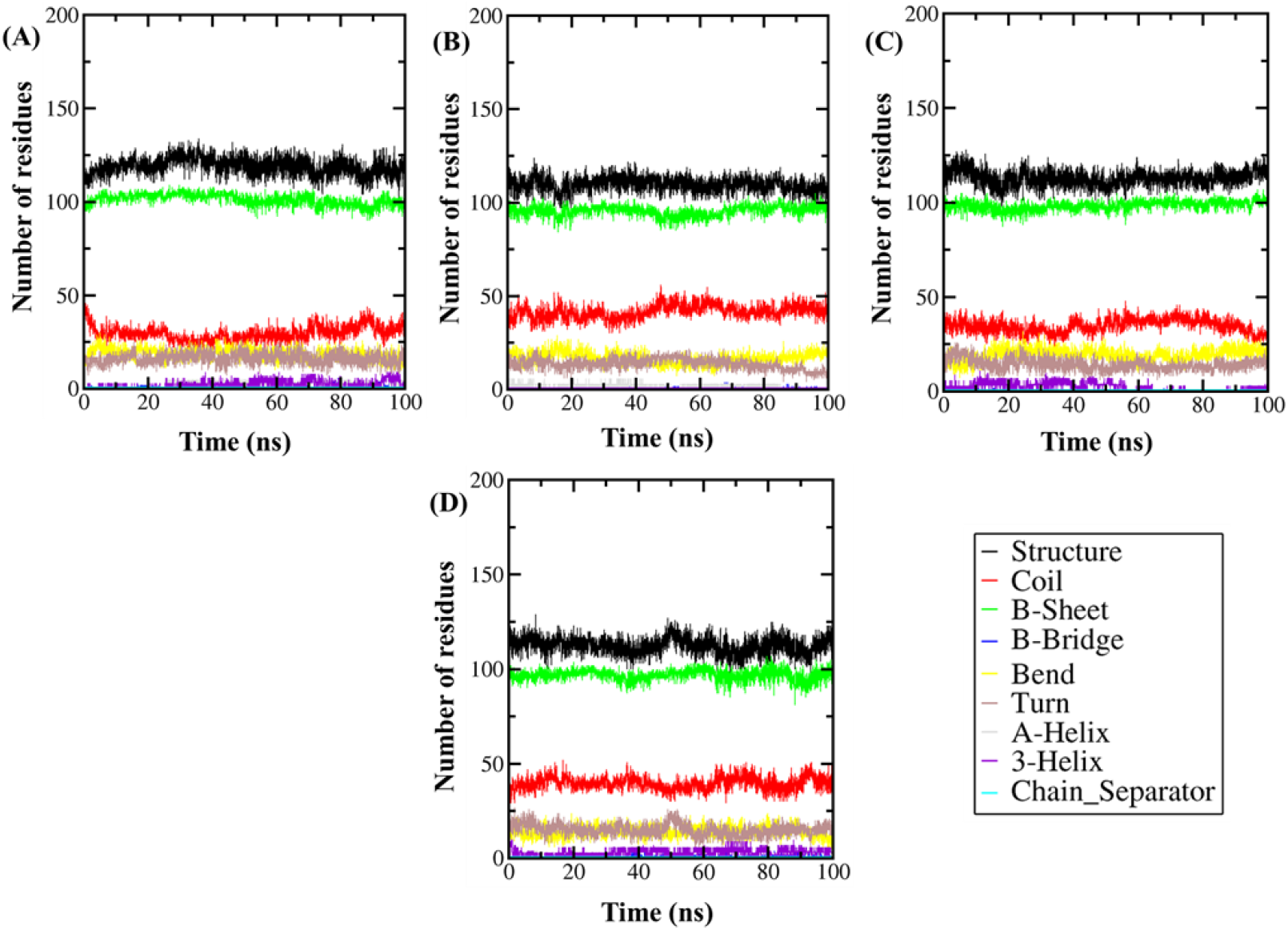
Secondary structure analysis of (A) SF, (B) SF-Doxycycline complex, (C) SF-Minocycline complex, (D) SF-Tetracycline complex. Structure = a-helix + b-sheet + b-bridge + Turn.

#### 3.2.6 Hydrogen Bond Analysis

The hydrogen bonding interactions between SF and the three ligands, Doxycycline, Minocycline, and Tetracycline, were analyzed over the 100 ns MD simulation were depicted in Figure 7. The *gmx hbond* tool was used with a donor–acceptor distance cutoff of 3.5 A° and an angle cutoff of 120° to calculate the mean number of hydrogen bonds formed during the simulation. The average number of hydrogen bonds formed by Doxycycline exhibited the highest average number of hydrogen bonds, maintaining 2–3 hydrogen bonds throughout the simulation, indicating stable interactions. Tetracycline also formed an average of two hydrogen bonds, but the interaction pattern was more variable and less consistent across the simulation period. Minocycline displayed the lowest average number of hydrogen bonds, forming on average one hydrogen bond with SF, but it was relatively stable throughout the simulation

**Figure 7.**
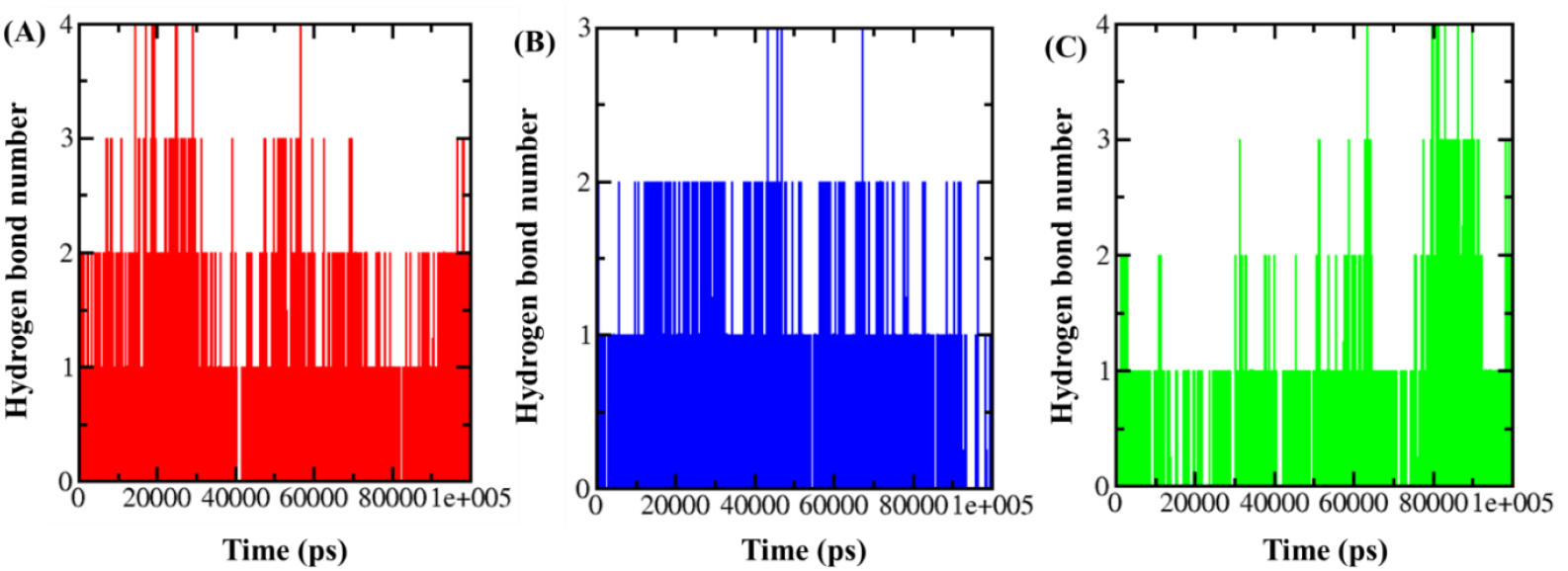
The average number of intermolecular hydrogen bonds that existed between Silk fibroin protein and (A) Doxycycline, (B) Minocycline, and (C) Tetracycline

### 3.3 MMPBSA analysis

Binding free energy calculations were performed to predict the average binding energies of the protein–ligand complexes and to gain insights into the energetic contributions of interacting residues. The energy parameters, including van der Waals, electrostatic, polar solvation, SASA, and total binding energy, are presented in Table 2. Doxycycline showed the strongest binding affinity with a total binding energy of –25.71 kcal/mol, indicating its potential as a stable binder to the silk fibroin (SF) matrix. This high affinity was primarily driven by favorable van der Waals interactions (–41.46 kcal/mol) and electrostatic energy (–11.23 kcal/mol), which outweighed the unfavorable polar solvation contribution (+30.92 kcal/mol). Tetracycline exhibited a binding energy of 18.27 kcal/mol, while Minocycline showed the least binding energy (–10.80 kcal/mol) among the three tested ligands. For all complexes, van der Waals energy emerged as the dominant stabilizing interaction, suggesting its central role in ligand binding within the SF environment. Overall, from the MMPBSA analysis, it is evident that Doxycycline interacts with SF much more strongly than Tetracycline and Minocycline.

**Table 2.**
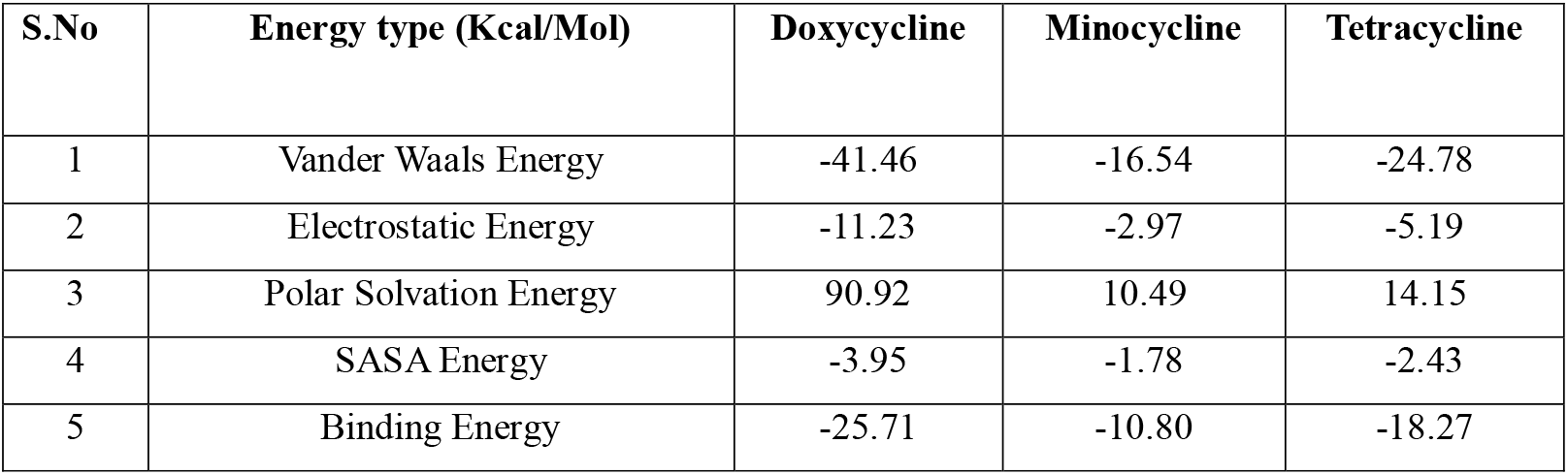
Table showing the MMPBSA-based estimation of the relative binding energies of different SF-ligand complexes.

| S.No | Energy type (Kcal/Mol) | Doxycycline | Minocycline | Tetracycline |
| --- | --- | --- | --- | --- |
| 1 | Vander Waals Energy | -41.46 | -16.54 | -24.78 |
| 2 | Electrostatic Energy | -11.23 | -2.97 | -5.19 |
| 3 | Polar Solvation Energy | 90.92 | 10.49 | 14.15 |
| 4 | SASA Energy | -3.95 | -1.78 | -2.43 |
| 5 | Binding Energy | -25.71 | -10.80 | -18.27 |

### 3.4 Superimposition analysis

The superimposition of SF protein and protein ligand complexes before and after simulation was performed using match matchmaker tool of UCSF Chimera to identify the structural changes protein and to monitor the orientation of the ligands following a 100 ns simulation. The alignment score of UCSF chimera quantifies the quality of superimposition between two structures by incorporating the number of matched residues and structural similarity. The illustrations of superimposed structures of SF and SF-ligand complexes before and after simulations are depicted in Figure 8.

**Figure 8.**
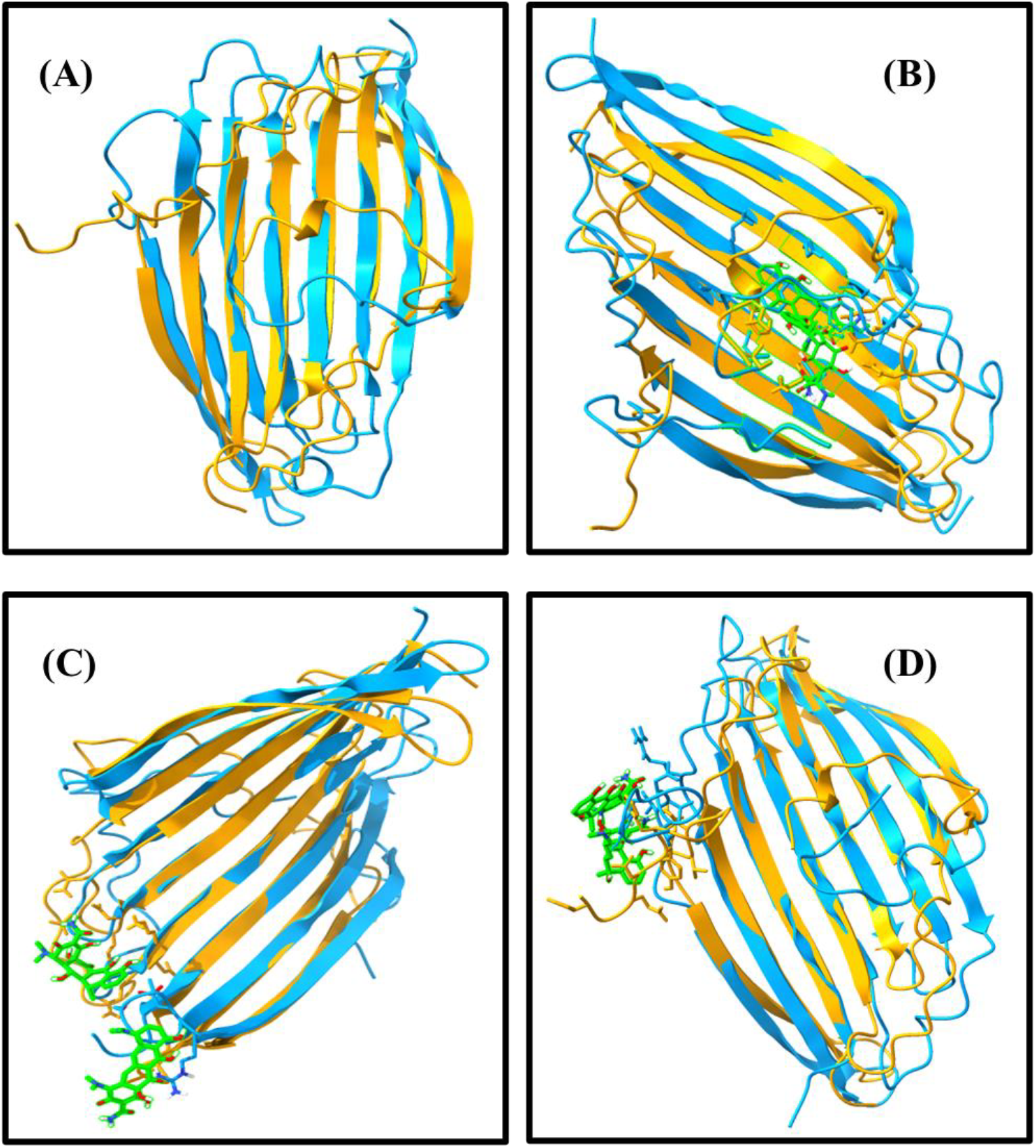
Superimposed structures of SF and SF-ligand complexes before and after 100 ns MD simulation. (A) SF (B), SF-Doxycycline complex (C), SF-Minocycline complex (D), SF-Tetracycline complex. All the structures before the simulation were presented in orange, and structures after the 100 ns simulation were represented in blue. The ligands were presented in green colour before and after simulation.

The SF after simulation displayed slight alterations in its beta sheet structure and drifting away from its original axis. Overall, the protein structure was stable and showed an acceptable RMSD (below 2.0 Å) score of 1.26 Å and an alignment score of 391.8. The SF-Doxycycline complex after simulation showed alterations at the site of interaction, and ligand orientation was stable, and no drift from the original site of interaction. The overall structure was stable and showed an RMSD score of 1.29 Å with an alignment score of 404.6. The SF-Minocycline complex also showed minimal alterations in the protein structure with an RMSD score of 1.194 Å and an alignment score of 409.2, but the orientation of the ligand was changed from the initial binding site, justifying the weaker interactions between minocycline and SF as validated by MMBPSA analysis. The SF-Tetracycline complex structure was stable with an RMSD score of 1.029 Å and an alignment score of 412.8.

## 4.0 Conclusion

The integrated approach of molecular docking and MD simulations demonstrated that Doxycycline exhibits the strongest and most stable interaction with SF when compared to Minocycline and Tetracycline. Among the three ligands, Doxycycline achieved the highest docking score and the most favourable binding free energy as determined by MM-PBSA analysis. In contrast, Minocycline showed a slightly lower docking score and the least favourable binding free energy. Although minocycline displayed a relatively stable RMSD and RMSF profile during MD simulations, Doxycycline consistently formed more stable interactions with SF. MD simulations conducted over 100 ns confirmed that the SF structure remained stable when complexed with Doxycycline, Minocycline, and Tetracycline, with only minor conformational fluctuations observed. These findings suggest that Doxycycline-loaded SF complexes offer promising potential for sustained and targeted antibiotic delivery. Nonetheless, since this study is limited to computational analyses, further in vitro and in vivo studies are required to validate the therapeutic efficacy of these complexes.

## Author Contributions

**Sreenivasan Kuravi:** Executing the experiments, Writing original draft, Methodology, software, analysis, and validation. **Kranti Kiran Reddy Ealla:** Conceptualisation, Formal analysis, Methodology, Project administration, Resources, Supervision. **Kiran Kumar Bokara**: Conceptualisation, Formal analysis, Methodology, Project administration, Resources, funding, Supervision. **Vikas Sahu:** Formal analysis, **Neema Kumari:** Formal analysis, writing-review. **Chandra Sri Durga**: Formal analysis and resources.

## Data availability

The data that support the findings of this study are available within the article

